# Modulation index-based phase–amplitude coupling does not encode temporal polarity

**DOI:** 10.64898/2026.06.05.730524

**Authors:** Mahmoud Keshavarzi

## Abstract

Phase–amplitude coupling (PAC) is widely used to quantify interactions between neural oscillations across timescales and is often interpreted as reflecting temporally meaningful coordination between slow and fast neural activity. Combining empirical EEG data with mathematical analysis, we show that the modulation index (MI), a widely used PAC metric, remains unchanged when the phase of the low-frequency oscillation is inverted by 180°. By contrast, preferred phase rotates by exactly 180°, as expected. Thus, opposite temporal organisations of cross-frequency coupling can yield identical MI values. This finding defines a fundamental interpretational limit of MI-based PAC: MI quantifies the strength of phase-dependent amplitude modulation, but not its temporal polarity. Accordingly, MI alone cannot distinguish between temporally aligned and temporally inverted coupling configurations or support inferences about temporal alignment, phase polarity, or directionality. Because PAC is widely used across neuroscience, these results establish an important boundary on what one of the field’s most common cross-frequency coupling measures can validly reveal about neural coordination.

## Introduction

Cross-frequency coupling, particularly phase–amplitude coupling (PAC), is widely used to characterise hierarchical interactions between neural oscillations operating at different timescales. More specifically, PAC is often interpreted as reflecting the organisation of higher-frequency activity by the phase of slower rhythms. In auditory neuroscience, for example, low-frequency oscillations such as delta and theta have been proposed to provide a temporal scaffold for higher-frequency activity in bands such as beta and gamma, thereby supporting the segmentation and encoding of speech and linguistic information [1]. Within this framework, the phase of slow oscillations is thought to regulate fluctuations in neuronal excitability and thereby influence the timing and amplitude of higher-frequency neural activity.

PAC is most commonly quantified using the modulation index (MI), which captures the extent to which high-frequency amplitude varies systematically as a function of low-frequency phase [2]. This approach builds on earlier demonstrations of cross-frequency coupling in human cortex [3] and has since been applied in studies of perceptual selection and attention [4–6], speech and language processing [7, 8], and working memory [9]. Crucially, PAC is often interpreted as reflecting temporally meaningful coordination between oscillations operating at different timescales.

Despite its widespread use, a key assumption underlying PAC analyses remains largely untested: that the phase relationship between slow and fast oscillations carries temporally meaningful information. This assumption is especially prominent in domains in which low-frequency phase is interpreted as shaping the timing of higher-frequency activity. In speech and language processing, for example, delta- and theta-band phase have been linked to prosodic and syllabic structure, respectively, and their temporal alignment is thought to shape the neural encoding of continuous speech [1, 10]. Here, combining empirical EEG data with mathematical analysis, we directly tested whether modulation index-based PAC distinguishes between opposite temporal organisations of cross-frequency coupling.

## Results

We directly tested this question using empirical EEG data recorded during continuous speech listening (Fig. 1a–h). Data were obtained from an open-access dataset ([11]; see Materials and Methods), and PAC was computed between delta (1–4 Hz) and beta (13–25 Hz), as well as between theta (4–8 Hz) and gamma (25–40 Hz). The modulation index was calculated using a standard phase-binning approach [2]. Critically, we inverted the phase of the low-frequency signal by adding π radians (180°), thereby imposing a half-cycle phase shift that reversed the temporal alignment between low-frequency phase and high-frequency amplitude while preserving spectral content.

**Figure 1.**
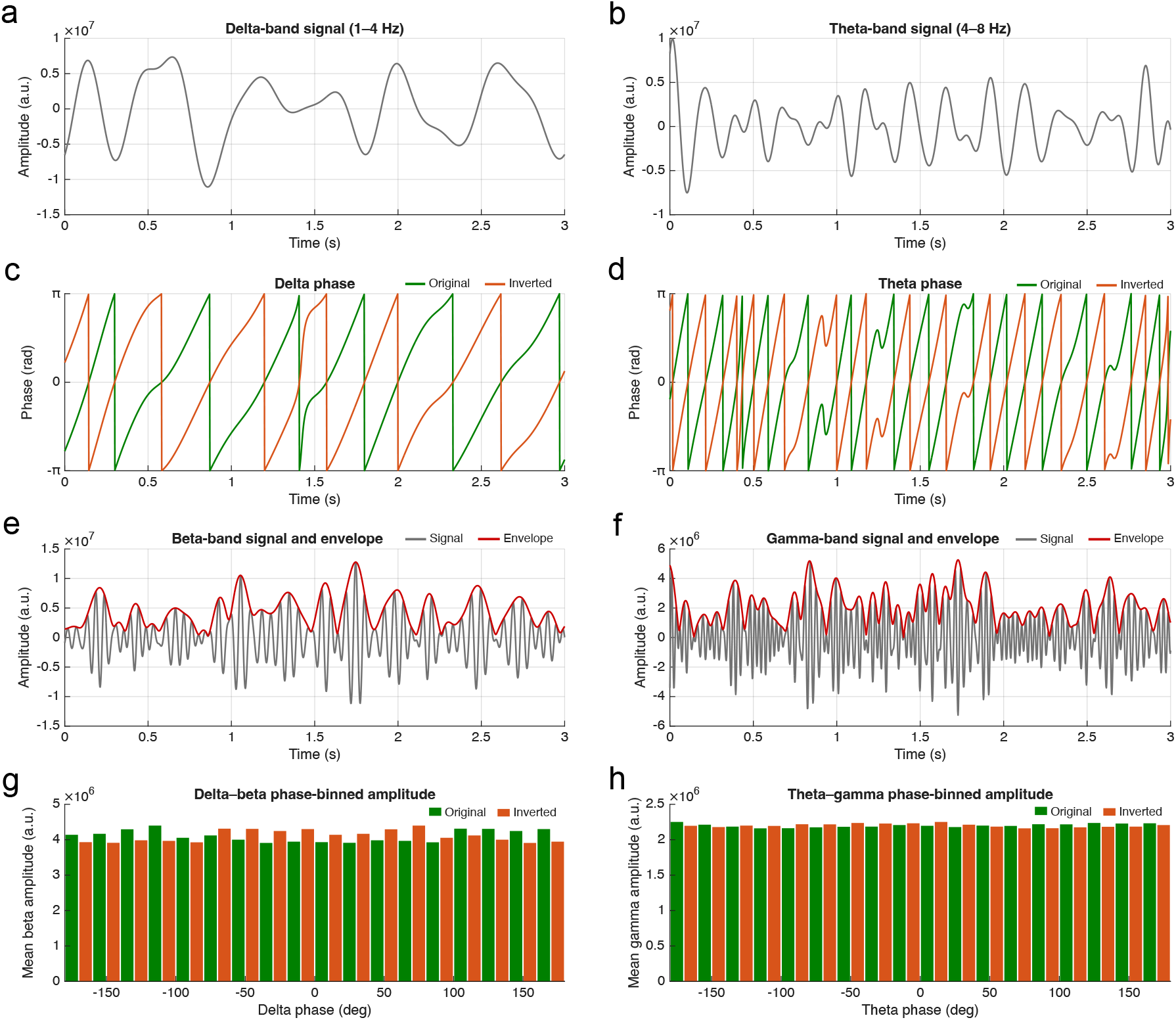
Phase inversion preserves phase-binned amplitude structure while shifting low-frequency phase by half a cycle. Representative EEG data from a single channel (A10) during continuous speech listening, shown for a 3-s segment. (a) Delta-band signal (1–4 Hz) extracted by band-pass filtering. (b) Theta-band signal (4–8 Hz) from the same segment. (c) Instantaneous phase of the delta signal (green) and its inverted version (orange), showing a 180° phase shift with circular wrapping. (d) Instantaneous phase of the theta signal and its inverted version. (e) Beta-band signal (13–25 Hz; grey) and its amplitude envelope (red), derived using the Hilbert transform. (f) Gamma-band signal (25–40 Hz; grey) and its amplitude envelope (red). (g) Mean beta-band amplitude plotted as a function of delta phase for the original (green) and inverted (orange) signals across 18 phase bins, showing identical modulation index (MI) values. (h) Mean gamma-band amplitude plotted as a function of theta phase for the original (green) and inverted (orange) signals, showing identical MI values.

Phase inversion imposed a half-cycle shift in the low-frequency rhythm, reversing its phase alignment with high-frequency amplitude. Yet the modulation index was unchanged in both analyses (delta–beta: MI = 2.7135 × 10^−4^ for both original and inverted signals; theta–gamma: MI = 2.2516 × 10^−5^ for both conditions; Fig. 2a,b). By contrast, the preferred phase rotated by exactly 180°, shifting from −110° to 70° in the delta–beta analysis and from −170° to 10° in the theta–gamma analysis (Fig. 2c,d). These findings show that preferred phase is sensitive to phase inversion, whereas modulation index-based PAC strength is not.

**Figure 2.**
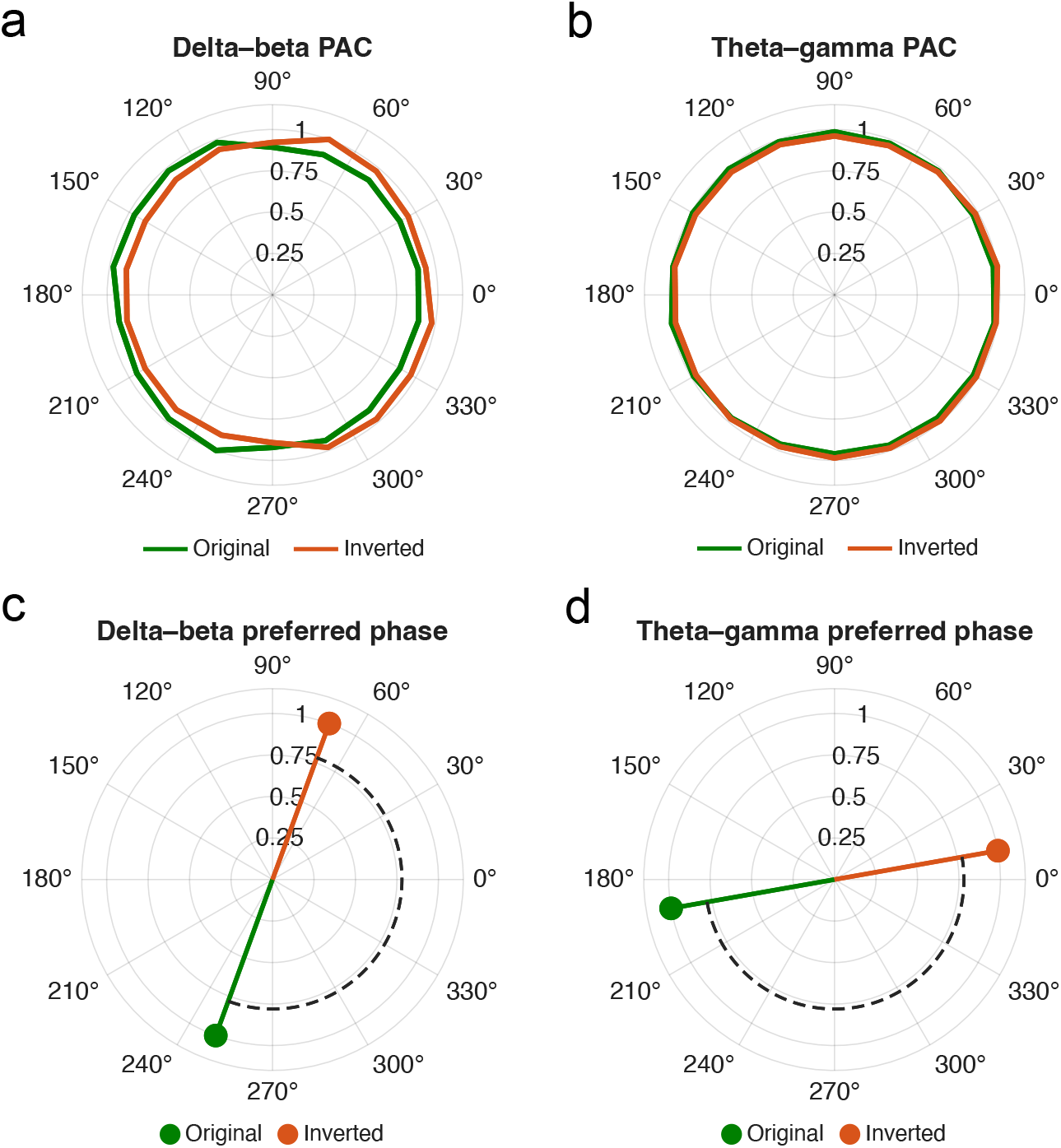
Modulation index is invariant to phase inversion, whereas preferred phase rotates by 180°. Polar plots of phase-binned high-frequency amplitude for (a) delta–beta and (b) theta–gamma PAC, respectively, shown for the original (green) and phase-inverted (orange) low-frequency signals. The phase-binned amplitude profiles remain highly similar across conditions, yielding identical modulation index (MI) values (delta–beta: 2.7135 × 10^−4^; theta– gamma: 2.2516 × 10^−5^). Preferred phase, defined as the phase bin associated with maximal high-frequency amplitude, is shown for (c) delta–beta and (d) theta–gamma PAC. Dashed lines indicate the angular separation between original and inverted conditions, showing an exact 180° rotation after phase inversion.

This invariance arises directly from the mathematical formulation of the modulation index. Because MI quantifies the deviation of the phase–binned amplitude distribution from uniformity [2], it is insensitive to circular shifts of the phase axis. A phase inversion corresponds to a rotation of the phase distribution by π radians, preserving the shape of the amplitude–phase relationship while altering its temporal alignment. Consequently, two signals with opposite temporal organisation produce identical PAC values.

## Discussion

Using empirical EEG data and mathematical analysis, we show that modulation index-based phase–amplitude coupling is invariant to 180° phase inversion of the low-frequency signal, even though preferred phase shifts by 180°. This finding has important implications for interpreting PAC across neuroscience, particularly in contexts where low-frequency phase is assumed to carry temporally meaningful information. In oscillatory models of speech processing, for example, low-frequency phase is proposed to align with linguistic structure in order to shape the timing of higher-frequency activity [1]. However, our results demonstrate that PAC magnitude alone cannot distinguish between correctly aligned and temporally inverted dynamics. Identical coupling strength can therefore arise from fundamentally different temporal organisations.

Our results extend previous methodological critiques of PAC, which have emphasized potential confounds and interpretational limitations [12, 13]. Although PAC robustly captures statistical dependencies between signals, it does not encode temporal directionality or phase polarity. This limitation is especially relevant in neurolinguistics, where theories often depend on the temporal alignment between neural activity and speech input.

In summary, we show that the modulation index is invariant to 180° phase inversion of the low-frequency signal. This fundamental property challenges the interpretation of modulation index-based PAC as evidence, on its own, for temporally meaningful neural coordination and indicates that PAC strength alone cannot support inferences about temporal alignment, phase polarity, or directionality. Accordingly, studies seeking to make mechanistic claims about temporal organisation should complement MI with measures that retain phase information, such as preferred phase or the full phase-binned amplitude profile.

## Materials and Methods

### Data and preprocessing

EEG data were obtained from an open-access dataset ([11], OpenNeuro ds004408). Recordings were acquired during continuous speech listening at 512 Hz using 128 scalp electrodes. Analyses were performed on continuous data. A representative channel (A10; channel 10) was selected for illustration, and all PAC analyses reported here were computed from this single-channel time series.

### Frequency decomposition

Signals were band-pass filtered using zero-phase FIR filters to extract delta (1–4 Hz), theta (4– 8 Hz), beta (13–25 Hz), and gamma (25–40 Hz) bands. For each filtered signal, the analytic signal was computed using the Hilbert transform. Instantaneous phase was extracted from the angle of the analytic signal for the delta and theta bands, whereas amplitude envelopes were extracted from the magnitude of the analytic signal for the beta and gamma bands.

### Phase–amplitude coupling

PAC was quantified using the modulation index [2]. Phase values were divided into 18 bins spanning −π to π. Mean high-frequency amplitude was computed within each phase bin and normalized to form a probability distribution. The modulation index reflects the divergence of this distribution from uniformity. PAC was computed separately for delta–beta and theta– gamma frequency pairs.

### Phase inversion and mathematical property of the modulation index

Phase inversion was implemented by adding π radians to the instantaneous phase of the low-frequency signal and wrapping the resulting values to the interval [−π, π). The invariance of the modulation index under this manipulation follows directly from its mathematical definition, as shown below. Following Tort et al. [2], the phase interval [−π, π) was divided into *N* uniformly spaced bins, and the mean high-frequency amplitude in bin *k* was denoted by *A*_*k*_, where *k* = 1, …, *N*. These values were normalised to form a probability distribution,

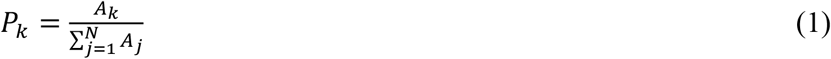

The modulation index (MI) was then defined as

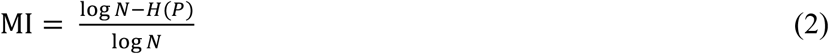

where *H*(*P*) is the Shannon entropy of the phase-binned amplitude distribution,

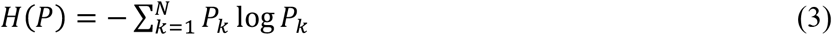

Now consider adding a constant phase offset, Δ*ϕ*, to the low-frequency phase, *ϕ*(*t*),

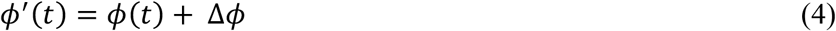

Exact invariance of the phase-binned MI holds whenever Δ*ϕ* is an integer multiple of the bin width 2*π*/*N*, that is,

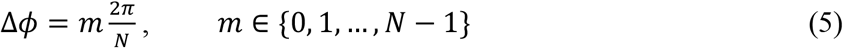

Here, *m* is the number of phase bins by which the phase distribution is shifted. Under such a shift, the phase bins are mapped exactly onto one another by a circular permutation. The phase-binned amplitudes and their normalised distribution therefore satisfy

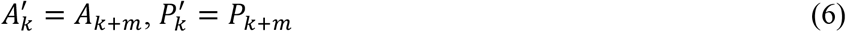

where indices are interpreted modulo *N*. Because Shannon entropy is invariant under permutation,

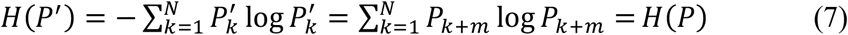

Substituting equation (7) into equation (2) gives

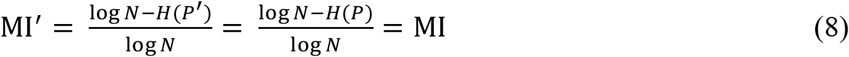

Therefore, the modulation index is invariant to any constant phase shift that maps the phase bins exactly onto one another. In the present analysis, *N* = 18, so phase inversion by *π* radians (180°) corresponds exactly to *π* = 9(2*π*/18), that is, a shift of 9 bins, and therefore leaves MI unchanged. By contrast, the preferred phase depends on the angular location of the bin with maximal high-frequency amplitude and therefore shifts with the applied phase rotation.

### Preferred phase

Preferred phase was defined as the phase bin associated with maximal high-frequency amplitude. It was identified separately for the original and inverted conditions, and circular differences were computed to quantify phase shifts.

## Data availability

The EEG data analysed in this study are publicly available from OpenNeuro (dataset ds004408). The MATLAB code used for the illustrative phase–amplitude coupling analyses has been made available to the journal during peer review in a private Code Ocean capsule and will be made publicly available upon publication.

